# Tissue Optimisation Strategies for High Quality Ex Vivo Diffusion Imaging

**DOI:** 10.1101/2021.12.13.472113

**Authors:** Rachel L. C. Barrett, Diana Cash, Camilla Simmons, Eugene Kim, Tobias C. Wood, Richard Stones, Anthony C. Vernon, Marco Catani, Flavio Dell’Acqua

## Abstract

*Ex vivo* diffusion imaging can be used to study healthy and pathological tissue microstructure in the rodent brain with microscopic resolution, providing a link between *in vivo* MRI and *ex vivo* microscopy techniques. A major challenge for the successful acquisition of *ex vivo* diffusion imaging data however are changes in the relaxivity and diffusivity of brain tissue following perfusion fixation. In this study we address this question by examining the combined effects of tissue preparation factors that influence image quality, including tissue rehydration time, fixative concentration and contrast agent concentration. We present an optimisation strategy combining these factors to manipulate the T1 and T2 of fixed tissue and maximise signal-to-noise ratio (SNR) efficiency. Applying this strategy in the rat brain resulted in a doubling of SNR and an increase in SNR per unit time by 135% in grey matter and 88% in white matter. This enabled the acquisition of excellent quality high-resolution (78 µm isotropic voxel size) diffusion data in less than 4 days, with a *b*-value of 4000 s/mm^2^, 30 diffusion directions and a field of view of 40 × 13 × 18 mm, using a 9.4 Tesla scanner with a standard 39 mm volume coil and a 660 mT/m 114 mm gradient insert. It was also possible to achieve comparable data quality for a standard resolution (150 µm) diffusion dataset in 2^1^/_4_ hours. In conclusion, the optimisation strategy presented here may be used to improve signal quality, increase spatial resolution and/or allow faster acquisitions in preclinical *ex vivo* diffusion MRI experiments.

## 1. Introduction

High-resolution *ex vivo* diffusion MRI offers unique opportunities to study tissue microstructure and the three-dimensional organisation of fibres in the rodent brain in both healthy animals and models relevant for human brain disorders. Compared to *in vivo* diffusion imaging, *ex vivo* experiments have fewer constraints in terms of scan time, which enables higher spatial resolution to be achieved. Due to changes in brain tissue properties after death and fixation however, *ex vivo* MRI and diffusion imaging in particular have greater challenges in achieving a sufficient signal-to-noise ratio (SNR), as found previously in human (Pfefferbaum et al., 2004; Miller et al., 2011; Roebroeck et al., 2019) and animal (D’Arceuil et al., 2007; D’Arceuil & de Crespigny, 2007; Dyrby et al., 2011, 2018) experiments. In animal experiments it is possible to fix the tissue by perfusion, which eliminates the interval between death and fixation, minimising tissue degradation associated with autolysis (Shepherd et al., 2009a). In this study we focus on tissue preservation in the rat brain. The same approach could easily be adapted for other animal species using perfusion fixation.

Tissue fixation is necessary to prevent decomposition and preserve the structure of *ex vivo* brain tissue. Among the changes associated with fixed tissue are decreases in T1 and T2 relaxation time (Tovi & Ericsson, 1992; Blamire et al., 1999; Pfefferbaum et al., 2004; Yong Hing et al., 2005; Thelwall et al., 2006; Blezer et al., 2007; D’Arceuil et al., 2007; Shepherd et al., 2009b). These changes in tissue relaxation affect the SNR of spin echo sequences, as the MR signal increases with shorter T1 but decreases with shorter T2. While signal loss from T2 relaxation can be minimised by shortening the echo time (TE), this poses a challenge for diffusion imaging, as the TE must be long enough to include both diffusion encoding and the imaging readout.

Fixed tissue has lower diffusivity than living tissue, reported in previous studies in the range of ∼ 2 – 8 ×10^−4^ mm^2^s^−1^ in white matter (Sun et al., 2003, 2005; D’Arceuil et al., 2007; Miller et al., 2011). This necessitates greater diffusion-weighting to achieve adequate contrast for advanced diffusion modelling methods. For example, Dyrby et al. showed that a *b*-value of approximately 4000 s/mm^2^ is desirable for multi-fibre estimation using *ex vivo* diffusion methods (2011) compared with 2500 - 3000 s/mm^2^ *in vivo* (Alexander & Barker, 2005; White & Dale, 2009; Tournier et al., 2013). Higher *b*-values and larger acquisition matrices require longer TE, which leads to signal loss from T2 relaxation, further highlighting the need to preserve SNR for *ex vivo* diffusion imaging.

To preserve SNR in high-resolution imaging, trade-offs must be made between SNR, voxel size and scan time. For example, SNR can be increased by acquiring multiple repetitions for signal averaging or by lengthening the repetition time (TR) until sufficient T1 recovery is acheived, both of which increase the total scan time. This is a limiting factor in high-resolution applications where scan time is already increased by the need for a larger acquisition matrix. Furthermore, in diffusion imaging, scan time is especially critical because it is multiplied by the number of volumes needed to sample different diffusion directions. The minimum number of directions required for DTI is six, but for robust tensor estimation, advanced modelling based on HARDI and advanced tractography applications, 30 - 60 directions are often recommended (Jones, 2004; Alexander & Barker, 2005; Tournier et al., 2013; Dell’Acqua & Tournier, 2019). The time needed to acquire 30 or more diffusion directions using a high-resolution spin echo sequence with adequate SNR (e.g. SNR *>*10 in non-diffusion-weighted images) quickly becomes prohibitive unless a strategy is used to improve SNR with respect to scan time, or SNR efficiency.

Existing approaches for improving SNR efficiency in *ex vivo* diffusion MRI include the development of pulse sequences such as diffusion-weighted Steady-State Free Precession (SSFP; McNab et al., 2009; Miller et al., 2011, 2012) and Gradient and Spin Echo (GRASE; Aggarwal et al., 2010), or hardware optimisation (Johnson et al., 2012; Calabrese et al., 2013; Calabrese & Johnson, 2013). The approach we consider here is to optimise the tissue preparation, which can be done alone or in combination with other approaches to improve SNR efficiency. In this study we combined multiple tissue preparation steps to manipulate the T1 and T2 characteristics of fixed rat brain tissue and maximise SNR efficiency. Specifically, we tested three tissue preparation factors: fixative concentration, duration of tissue rehydration after fixation, and the addition of a gadolinium-based MR contrast agent.

First, we consider fixative concentration. Animal *ex vivo* experiments usually involve fixation by trans-cardiac perfusion with paraformaldehyde (PFA) at a concentration of 4% by volume. Previous studies have shown an association between higher concentrations of PFA or formaldehyde and shorter relaxation time, particularly for T2 (Fishbein et al., 2007; Shepherd et al., 2009b; Birkl et al., 2018). Cahill et al. (2012) experimented with PFA concentrations from 0.5% to 4% in small samples of human brain tissue and found that concentrations between 2 - 4% were optimal. We therefore tested whether a reduced fixative concentration of 2% PFA would have an effect on SNR efficiency.

The second tissue preparation factor we consider for its effect on relaxation times was the rehydration of fixed tissue in phosphate buffered saline (PBS). Previous studies show substantial increases to T2 as a result of rehydration with PBS (Shepherd et al., 2005, 2009b; Thelwall et al., 2006; Blezer et al., 2007; D’Arceuil et al., 2007; Leprince et al., 2015), with T1 increasing slightly (D’Arceuil et al., 2007; Leprince et al., 2015) or not at all (Shepherd et al., 2005, 2009b; Thelwall et al., 2006; Blezer et al., 2007), indicating that this could be a beneficial strategy for maximising signal. The specimen size and rehydration duration vary in the literature, so one of our aims was to quantify the changes in relaxation constants over time immersed in PBS after perfusion fixation and determine the optimal tissue rehydration time.

The final tissue preparation factor we tested was the addition of the gadolinium-based contrast agent Gd-DTPA (Magnevist). Gadolinium reduces T1 and, to a lesser extent, T2 relaxation time, which allows TR to be shortened without losing signal from longitudinal relaxation (Benveniste & Blackband, 2002; Johnson et al., 2002; D’Arceuil et al., 2007; Huang et al., 2009). D’Arceuil et al. (2007) use the relationship between the contrast agent concentration and relaxivity to optimise SNR efficiency for fixed macaque brains immersed in PBS with Gd-DTPA. Diffusion MRI results obtained from gadolinium-enhanced rodent brain tissue have previously been compared with axonal tracing (Calabrese et al., 2015) and histology (Calabrese & Johnson, 2013; Wang et al., 2019, 2020) to establish the validity of this technique with respect to diffusion MRI connectivity and metrics of microstructure. In this study we used the ‘active staining’ technique (Johnson et al., 2002; Calabrese et al., 2013), in which a high concentration of contrast agent is introduced during perfusion fixation, followed by a lower concentration during the rehydration stage. We then followed the approach used in D’Arceuil et al. (2007) of measuring relaxivity at different concentrations to determine the optimum amount of Gd-DTPA to be added to the perfusate. In addition to our main experiment using Gd-DTPA, we have included a comparison between Gd-DTPA and the more recently available gadobutrol (Gadovist).

After characterising the diffusion and relaxation properties of fixative concentration, rehydration time and contrast agent concentration in the rat brain, we modelled how these factors can be combined to maximise SNR efficiency and demonstrated the resulting improvement in SNR. We then present an application of tissue optimisation in high-resolution and more conventional diffusion acquisitions. Finally, we used histology data to verify the quality of fixation in our modified tissue preparation.

## 2. Materials and methods

### Animal preparation

All animal experiments were conducted with approval from the local King’s College London ethics committee in accordance with the UK Home Office Animals (Scientific Procedures) Act 1986. Adult male rats (Sprague Dawley, n = 30) were euthanised with pentobarbital (60 mg/kg i.p.) and transcardially perfused for approximately two minutes at 100 ml/min pump speed with a minimum of 200 ml ice-cold heparinised 0.9% saline (50 IU/ml) followed by a minimum of 200 ml of either 2% or 4% paraformaldehyde (PFA, ‘Parafix’, Pioneer Research Chemicals, UK) buffered at 7.4 pH containing Gd-DTPA in a concentration range of 0 - 50 mM. After perfusion, the heads were removed and immersed in the fixation mixture for four days. We chose a post-fixation period of four days to ensure adequate fixation when experimenting with the reduced fixative concentration, however it may be sufficient to post-fix for shorter periods, e.g. 24 hours or less, as done elsewhere in rats fixed with the standard fixative concentration (Johnson et al., 2012; Laitinen et al., 2015). The samples were then rehydrated in 50 ml phosphate buffered saline (PBS) with 0.05% sodium azide preservative and either 0 or 1 mM Gd-DTPA. Two samples were prepared similarly using gadobutrol instead of Gd-DTPA. Details of the fixative concentration, contrast agent concentration and rehydration time for all samples are summarised in Table 1. All samples were refrigerated at 4 °C during rehydration and moved to room temperature four hours before scanning. For the rehydration and gadolinium experiments, brains remained *in situ* to prevent potential tissue damage from skull-removal and handling, and excess tissue was removed from around the skull. For the high-resolution diffusion acquisition, the brain was removed from the skull prior to scanning to reduce the field of view in order to minimise scan time. For scanning, samples were sealed in plastic tubes padded with gauze to prevent movement and immersed in proton-free fluorinated liquid (Galden; Solvay) to reduce susceptibility artefacts.

**Table 1:** Rat preparation parameters for rehydration and gadolinium experiments, optimised high-resolution diffusion protocol and contrast agent comparison. n number of rats used for each experiment, CA contrast agent. ^*^Rats perfused with 35mM Gd-DTPA were rehydrated for 55 - 56 days.

| Experiment | *n* | [Fixative] | [CA] in perfusate | [CA] in rehydration solution | Rehydration interval (days) |
| --- | --- | --- | --- | --- | --- |
| Rehydration experiment | 4 | 4% (n=2), 2% (n=2) |  |  | 0, 2, 4, 6, 14, 24, 35 (scanned sequentially) |
| Gadolinium experiment | 20 | 4%, 2% (n=2 for each [CA]) | 0, 15, 25, 35, 50 mM Gd-DTPA | 1 mM Gd-DTPA | 35-38* |
| Reference (no gadolinium) | 2 | 4% (n=1), 2% (n=1) |  |  | 35 |
| High- and mid-resolution diffusion | 1 | 2% | 15 mM Gd-DTPA | 1 mM Gd-DTPA | $\leq 35$ |
| Contrast agent comparison | 4 | 4%, 2% (n=2 for each [CA]) | 5, 15 mM gadobutrol | 1 mM gadobutrol | 35 |

### MRI hardware

MR imaging was performed at the BRAIN Centre (brain-imaging.org), King’s College London on a 9.4T Bruker Biospec scanner with a 39 mm volume coil (Rapid Biomedical GmbH, GER). The rehydration, gadolinium and contrast agent comparison experiments, and the mid-resolution diffusion acquisition were carried out with a 660 mT/m 114 mm gradient set. The high-resolution diffusion acquisition was carried out using a 1000 mT/m 60 mm gradient insert.

### Relaxometry

T1 maps were acquired using a 2D Rapid Acquisition with Relaxation Enhancement (RARE) sequence with variable TR = 200, 400, 800, 1500, 3000 and 5500 ms, TE = 7 ms, RARE factor = 2, acquisition matrix = 128 × 128 × 7 and voxel size = 0.22 × 0.22 × 1.00 mm. T2 maps were acquired using a multi-slice multi-echo sequence with TR = 2000 ms, TE = 8, 16, 24, 32, 40, 48, 56, 64, 72, 80, 88, 96, 104, 112, 120, 128, 136, 144, 152, 160, 168 and 176 ms, and the same geometric parameters as used for T1 mapping. T1 and T2 relaxation parameters where estimated using non-linear least squares curve fitting in MATLAB (version 9.6.0, MathWorks). The average T1 and T2 within regions of interest in the corpus callosum and cortex were calculated to compare samples (Fig. 1). For the rehydration experiment, T1 and T2 values were plotted against rehydration time.

**Figure 1:**
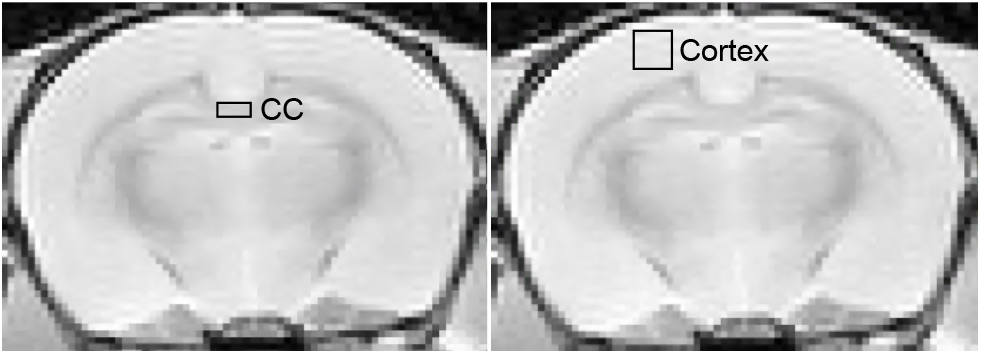
Regions of interest in the corpus callosum (CC) and cortex used in the rehydration and gadolinium experiments.

For the gadolinium experiment, T1 and T2 were modelled with respect to the concentration of contrast agent according to the relationship,

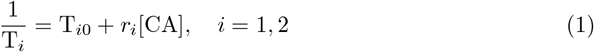

(Lauffer, 1987), where T_*i*_ is the observed relaxation time; T_*i*0_ is the baseline relaxation time, which in this case represents the tissue soaked in PBS with 1mM of contrast agent; *r*_*i*_ is the relaxivity of the contrast agent in the tissue; and [CA] refers to the concentration of contrast agent added to the perfusate during fixation.

### DTI measurements for optimisation

For the rehydration and gadolinium experiments, diffusion tensor imaging (DTI) measurements were carried out using a 2D diffusion weighted Echo Planar Imaging (EPI) sequence with a target *b*-value of 1500 s/mm^2^, 30 diffusion- weighted and 5 non-diffusion-weighted images, TR = 1000 ms, TE = 21 ms, 4 segments, acquisition matrix = 128 × 128 × 7 and voxel size = 0.23 × 0.23 × 1.00 mm. The data were denoised using Marchenko-Pasteur principal component analysis noise estimation for diffusion MRI Veraart et al. (2016) and Gibbs-ringing correction as described in Kellner et al. (2016). The data were then corrected for eddy current distortions and diffusion tensor parameters were estimated using ExploreDTI 4.8.6 (www.exploredti.com). Fractional anisotropy (FA) and mean diffusivity (MD) were recorded for regions of interest in the corpus callosum and plotted against rehydration time or [Gd-DTPA].

### SNR efficiency optimisation

SNR efficiency was calculated in terms of T1, T2, TR and TE, where T1 and T2 were modelled in terms of [Gd-DTPA] as described above using Equation 1. SNR efficiency is defined as 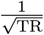 SNR and given by the following formula:

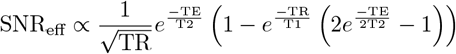

D’Arceuil et al. (2007). This allowed us to predict SNR efficiency for any given combination of [Gd-DTPA], TR and TE.

### SNR comparison

To compare SNR between standard and optimised tissue preparations, one rat brain fixed with 4% fixative and no gadolinium, rehydrated in PBS, was compared with another fixed with 2% fixative and 15 mM Gd-DTPA, rehydrated in PBS with 1 mM Gd-DTPA. A 3D diffusion-weighted spin echo acquisition was used with one diffusion- weighted volume (*b* = 2500 s/mm^2^) and one non-diffusion-weighted volume, with TR = 250 ms, TE = 26.82 ms, flip angle = 90°, acquisition matrix = 104 × 93 × 8 and isotropic voxel size = 0.25 mm, and scan time of 6.2 minutes.

Mean signal (*η*) values were recorded for regions of interest in the corpus callosum and cortex, while the standard deviation of the noise (*σ*_noise_) was measured from a region outside the sample. SNR was then estimated using the formula from Jones & Basser (2004):

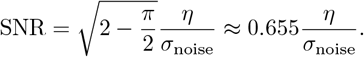

### High and mid-resolution diffusion protocols

Two examples of diffusion protocols were acquired using the current optimised tissue preparation approach, one high and one midresolution protocol. The high-resolution example was acquired using a 3D diffusion-weighted spin echo sequence with TR = 250 ms, TE = 26.78 ms, *b* = 4000 s/mm^2^, *δ* = 3 ms, Δ = 11 ms, maximum gradient amplitude = 577 mT/m, flip angle = 90°, 30 diffusion directions and 3 non-diffusion-weighted volumes. The isotropic voxel size was 78 µm, acquisition matrix = 512 × 162 × 232 and total scan time was 91.35 hours (2.77 hours per volume).

The mid-resolution example diffusion protocol was acquired using 3D diffusion-weighted EPI with TR = 280 ms, TE = 26.16 ms, *b* = 2000 s/mm^2^, *δ* = 4 ms Δ = 12.5 ms, gradient amplitude 405 mT/m, flip angle = 90°, 8 segments, 4 averages, 30 diffusion and 5 non-diffusion-weighted images, isotropic voxel size = 150 µm, acquisition matrix = 108 × 80 × 108, and scan time = 9.13 hours (16.6 minutes per volume). The EPI data was processed as described in *DTI measurements for optimisation*

The high-resolution spin echo dataset was denoised as described above. As spin echo data is robust to eddy current and geometric distortions, no further corrections were required. Diffusion modelling and tractography processing were done using StarTrack (www.natbrainlab.co.uk). Whole brain tractography data was generated using both DTI and spherical deconvolution models. For DTI tractography, a stopping threshold of FA = 0.1, angle threshold 45° and step size 78 µm were used. For spherical deconvolution, the fibre orientation distribution function was estimated using the damped Richardson-Lucy algorithm described in Dell’Acqua et al. (2010) with parameters *α* = 0.6, *n* = 0.001, *r* = 8 and 500 iterations. The tractography stopping threshold for this model was HMOA = 0.002, angle 45° and step size 78 µm, as above. Tracts were dissected in Trackvis 6.0.1 (www.trackvis.org) using manually drawn regions of interest with inclusion and exclusion conditions.

### Histology

Histology and immunohistochemistry were performed to test whether reducing the fixative concentration or adding Gd-DTPA to the fixation and rehydration stage had an impact on the quality of the fixed tissue. Three samples were compared, one fixed with 4% PFA, no contrast agent, i.e. the standard protocol, one fixed with 2% PFA, no contrast agent to test the effect of reduced fixative concentration, and one sample fixed with 2% PFA and 15 mM Gd-DTPA to test the effect of active staining with a gadolinium contrast agent. Once scanning had been completed, the brains were extracted from the skulls, cryoprotected in 30% sucrose and sectioned at 35 µm thickness. Tissue sections were stained with Cresyl Violet for Nissl bodies in neurons, Luxol Fast Blue for myelin, or IBA- 1 antibody using immunohistochemistry testing for microglia. Slides from all three stains were scanned with an Olympus VS120 slide scanner at ×40 magnification. The results were compared by inspection of expected cell and tissue morphologies, according to experience of previously immuno-stained tissues using the same protocols.

## 3. Results

### Rehydration experiment

The first tissue preparation factor we investigated was the rehydration of fixed rat brains in PBS prior to scanning. We found that rehydration resulted in increased T2 following an exponential recovery curve with relatively little to no effect on either T1 or diffusion metrics. Figure 2 shows how these variables change over 35 days of rehydration in a single volume of PBS. For the 4% fixative model, T1 was relatively stable, changing only from 1576 to 1618 ms (3% increase) in the white matter of the corpus callosum and 1631 to 1724 ms (5% increase) in the cortex after 35 days rehydration. The 2% fixative model had similar initial values of T1 that increased slightly over the 35-day period, from 1659 to 1781 ms (7% increase) in white matter and 1742 to 1781 ms (2% increase) in grey matter. A greater effect was seen in the T2 results. For the 4% fixative model, the T2 was initially 32 ms and increased to 55 ms (72%) in white matter, and from 33 to 60 ms (45% increase) in grey matter. For the 2% fixative model, T2 increased from 41 to 61 ms (50%) in white matter and 44 to 58 ms (24%) in grey matter. T2 stabilised faster for the 2% fixative model than the 4% model, taking 17 days for white and 14 days for grey matter to increase to within 95% of the model plateau value, compared to 32 days for white and 47 days for grey matter in the 4% model. Following from this result, the subsequent experiments were conducted on samples rehydrated for 35 days to ensure that T2 was fully recovered in the white matter for both the 2% and 4% fixative models, assuming that the rehydration time-curves are similar for samples in PBS with and without contrast agent.

**Figure 2:**
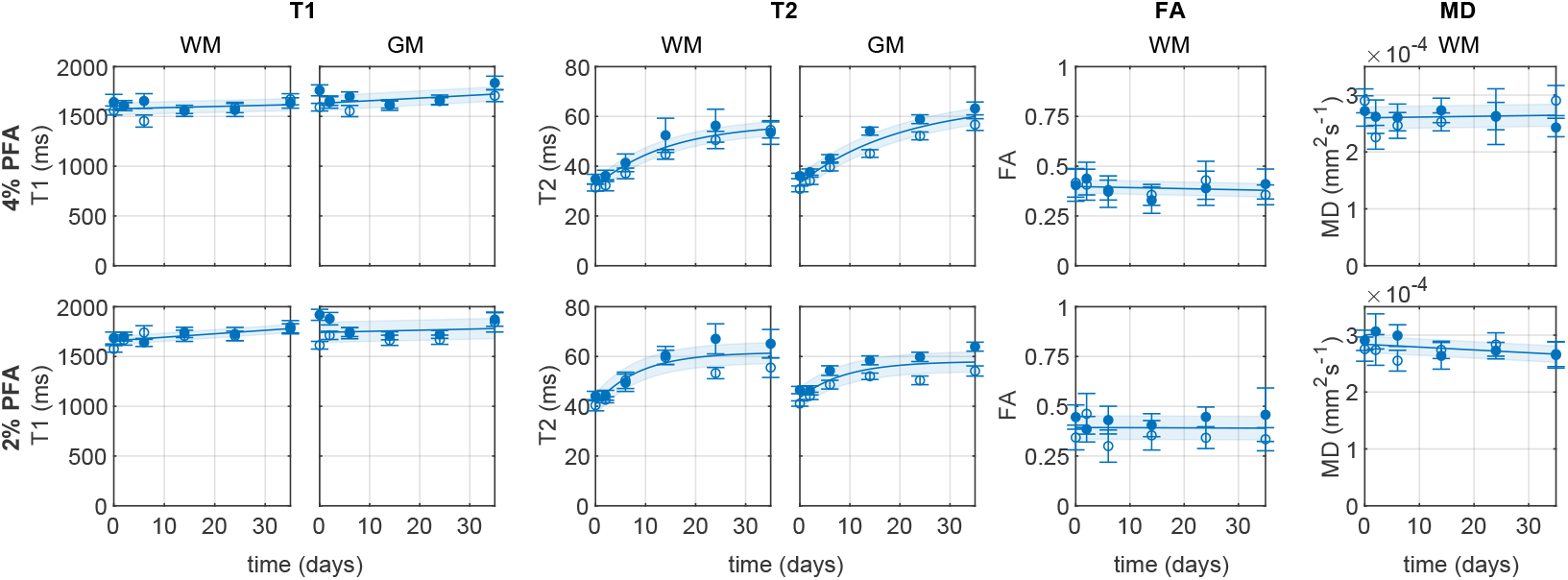
Plots showing how T1, T2, FA and MD vary with number of days soaking in PBS for rat heads fixed with 4% (top row) and 2% (bottom row) fixative. Data points show mean and standard deviation values within a region of interest in the white matter (WM) of the corpus callosum, and cortical grey matter (GM) for a single rat brain. Lines of best fit *±* standard error of the regression are shown.

Diffusion metrics FA and MD showed slight fluctuations in the first week of rehydration but changed minimally overall during the rehydration period. FA changed from 0.40 to 0.38 over the 35 days for the 4% fixative model and remained at 0.39 for the 2% fixative model. MD changed from 2.60 × 10^−4^ to 2.65 × 10^−4^ mm^2^s^−1^ after 35 days for the 4% fixative model, and from 2.84 to 2.67 × 10^−4^ mm^2^s^−1^ for 2% fixative.

### Gadolinium experiment

The following experiment was conducted to enable us to further manipulate T1 and T2 by introducing the gadolinium-based contrast agent Gd-DTPA. Figure 3 shows the relationship of T1, T2 and diffusion properties to the concentration of Gd-DTPA used in fixation. As expected, both T1 and T2 decrease with an inverse relationship to Gd-DTPA concentration according to the model in Equation 1, with T1 decreasing more rapidly than T2. The estimated relaxation parameters for the model are given in Table 2. Initial values for T1 and T2 represent the baseline case in which rats were rehydrated in 1 mM Gd-DTPA. T1 was initially 714 ms in white matter and 706 ms in grey matter for the 4% fixative model, and 882 ms in white and 896 ms in grey matter for the 2% fixative model. This represents a 23% increase in white and 27% in grey matter T1 from lowering the fixative concentration. At higher concentrations of Gd-DTPA greater than ∼10 mM, the T1 curves for the two models appeared to converge. Comparing the T2 results between the two fixative preparations, we also found higher initial values for the 2% fixative model at lower concentrations of contrast agent. T2 was initially 36 ms in white and 37 ms in grey matter for the 4% fixative model and 47 ms in white and grey matter in the 2% model, representing an increase of 31% in white and 27% in grey matter by lowering the fixative concentration. The T2 curves appeared to converge at ∼50 mM. The models for grey and white matter were very similar, with inter-individual differences and the effect of contrast agent being much greater than grey-white differences within rats.

**Table 2:** Parameters used to model relaxation time with respect to contrast agent concentration, according to Equation 1. T_*i*0_ is the baseline relaxation time for tissue fixed with no contrast agent and rehydrated in 1 mM contrast agent, units in milliseconds; *r*_*i*_, is the relaxivity constant for contrast agent added during fixation, units in mM^−1^s^−1^; and *σ*_*i*_ is the standard error of the regression for the model fitting, *i* = 1, 2, units in milliseconds. For Gd-DTPA, *n* = 10 for each fixative concentration. For the gadobutrol model, *n* = 4. Abbreviations: CA, contrast agent; WM, white matter; GM, grey matter.

| Fixation | CA | Tissue | $T_{10}$ | $r_{1[\text{CA}]}$ | $\sigma_1$ | $T_{20}$ | $r_{2[\text{CA}]}$ | $\sigma_2$ |
| --- | --- | --- | --- | --- | --- | --- | --- | --- |
| 4% PFA | Gd-DTPA | WM | 707 | 0.120 | 93.4 | 35.8 | 0.703 | 3.50 |
|  |  | GM | 706 | 0.149 | 83.6 | 36.9 | 0.709 | 4.13 |
| 2% PFA | Gd-DTPA | WM | 876 | 0.109 | 94.0 | 45.0 | 0.444 | 8.20 |
|  |  | GM | 896 | 0.167 | 90.6 | 46.8 | 0.801 | 7.75 |
| 2% PFA | gadobutrol | WM | 354 | 0.267 | 67.1 | 30.4 | 0.438 | 2.90 |
|  |  | GM | 385 | 0.325 | 21.7 | 31.2 | 0.516 | 1.80 |

**Figure 3:**
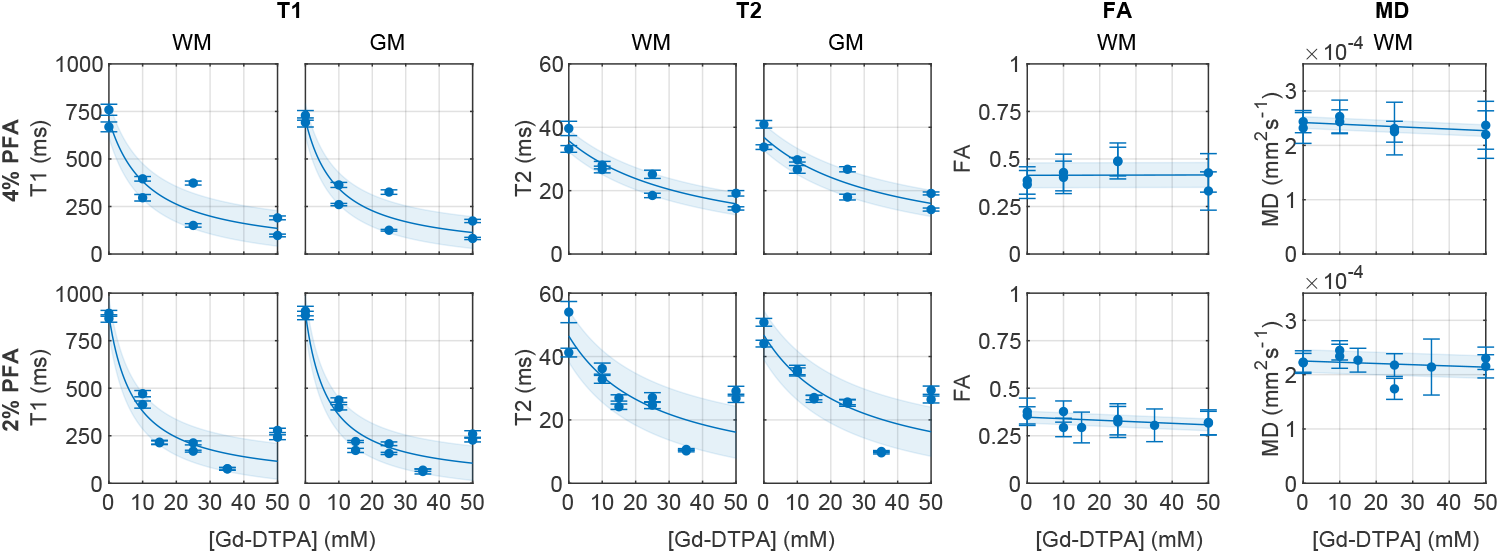
Plots showing how T1, T2, FA and MD vary with the concentration of Magnevist (Gd-DTPA) used in perfusion with 4% (top row) and 2% (bottom row) fixative. As above, data points and error bars represent the mean and standard deviation of values in the corpus callosum white matter (WM) and cortical grey matter (GM) for a single rat brain. T1 and T2 are modelled with Equation 1, FA and MD with lines of best fit. The haded region represents the standard error of the regression. All samples in this experiment, including those with no contrast agent in the perfusate, were rehydrated for at least 35 days in PBS with 1mM Gd-DTPA.

We also found that the addition of contrast agent had relatively little effect on diffusion metrics. FA in the 4% fixative model changed from 0.41 with no contrast agent in the fixative to 0.42 with 50mM Gd-DTPA, and in the 2% fixative model from 0.35 to 0.31. MD in 4% model changed from 2.42 × 10^−4^ to 2.27 × 10^−4^ mm^2^s^−1^, and in the 2% model from 2.25 × 10^−4^ to 2.14 × 10^−4^ mm^2^s^−1^.

In addition to the experiment above using Gd-DTPA, we included an analysis of the contrast agent gadobutrol in rats fixed with 2% fixative for comparison (Fig. 4). We found that the observed T1 in the gadobutrol model was shorter than in the Gd-DTPA model for equivalent concentrations by a factor of ∼ 0.4 in the white matter region and 0.4 − 0.5 in grey matter for the concentrations modelled. T2 in the gadobutrol model was shorter by a factor of 0.7 initially for both white and grey matter regions, while the difference between the two models reduced for higher concentrations. The estimated parameters for the gadobutrol model are given in Table 2.

**Figure 4:**
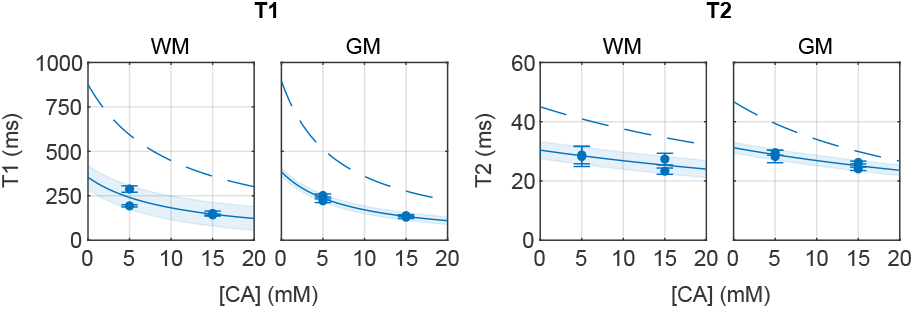
Comparison of relaxation time vs concentration curves for Gd-DTPA (dashed line) and gadobutrol (data points and solid line). Graphs show how T1 and T2 vary with concentration of contrast agent (CA) used in perfusion. A constant 1 mM CA is used in the rehydration stage for both models, including when the concentration in the perfusate is at 0 mM. The differing initial values of the graphs represents the effect of adding CA to the rehydration solution alone. Data points and error bars represent the mean and standard deviation of values in the corpus callosum white matter (WM) and the cortical grey matter (GM). Lines of best fit *±* standard error of the regression are shown.

The initial point of the graphs represents the case where 1 mM of each contrast agent was used in the rehydration stage only, with no contrast agent added during fixation. The difference between the two models here implies that the 1 mM of contrast agent used rehydration had a significant effect. It is worth noting that the concentration values shown on the *x* -axes in Figures 3 and 4 refer to the amount of contrast agent added to the perfusate, which may differ from the actual concentration in the tissue at the time of scanning. For reference, we also measured T1 and T2 in rats with no contrast agent in either the perfusate or the rehydration solution. In the 4% fixative model with no contrast agent, T1 = 1705 *±* 91 ms in white matter and 1760 *±* 181 ms in grey matter and T2 = 52 *±* 2 ms in white and 51 *±* 12 ms in grey matter, (*n* = 3). In the 2% fixative model, T1 = 1824 *±* 63 ms in white and 1828 *±* 81 ms in grey matter and T2 = 61 *±* 5 ms in white matter and 57 *±* 8 in grey matter (*n* = 3).

### SNR efficiency optimisation

After characterising the effects of different tissue preparation factors on T1 and T2, we aimed to find the concentrations of Gd-DTPA that would maximise SNR efficiency. This was done taking into consideration realistic constraints on TE and TR. Figure 5 shows how SNR efficiency varies with TR, TE and Gd-DTPA. First, we consider TE. In general, SNR efficiency is maximised when the TE is short, but in practice, the minimum available TE is limited by scanner capabilities or other experimental factors such as matrix size and diffusion weighting. Secondly, we consider the choice of TR. A longer TR can increase SNR but also increases the scan time, resulting in lower SNR efficiency after a certain point. Thirdly, we consider the concentration of Gd-DTPA, which can be varied freely to achieve optimal SNR efficiency for a given TE and TR. Figure 5A shows the SNR efficiency for a range of TE and TR resulting from an optimal choice of Gd-DTPA concentration. The optimal TR for a given TE is indicated in white. Figure 5B shows the concentrations of Gd-DTPA that lead to maximum SNR efficiency, corresponding to Figure 5A. We observe that more contrast agent is required for shorter TE and TR and vice versa. To illustrate how SNR efficiency varies with contrast agent concentration for a given TE and TR, Figure 5C shows the SNR efficiency profiles for 4 selected examples: TE/TR = 27/250 ms, ii) 27/800 ms, iii) 15/250 ms, iv) 15/800 ms. We observe that the shape of the SNR efficiency curves varies for the different TRs shown, with narrower peaks for the longer TR values, and the range of concentrations for optimising SNR efficiency is wider for shorter TR and narrower for longer TR. As a guide for selecting the contrast agent concentration, we calculated the concentration range of Gd-DTPA required to give within 5% of the maximum SNR efficiency, for the four example cases given in Figure 5. These are i) 12-31 mM, ii) 3-15 mM iii) 19-46 mM, and iv) 6-22 mM.

**Figure 5:**
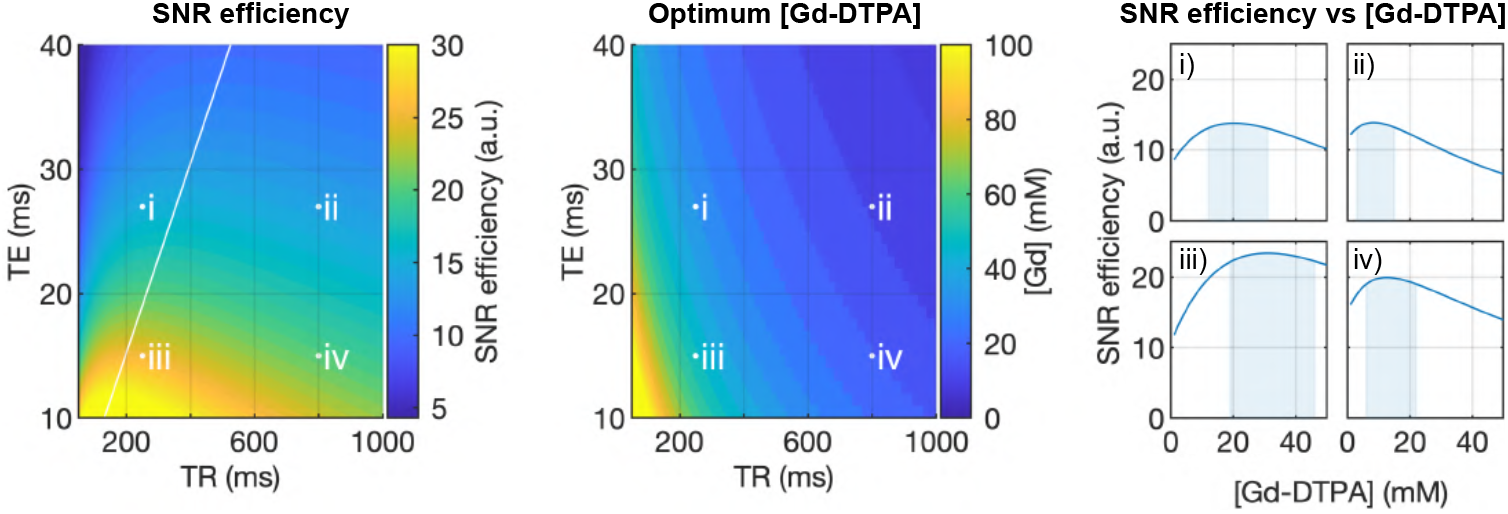
Plots showing how SNR efficiency varies with TE, TR and the concentration of Gd-DTPA used in fixation. Left) SNR efficiency as a function of TE and TR, given the optimum [Gd-DTPA] for each combination. This panel can be used to determine the optimal TR for a given TE, following the white line plotted on the graph. Middle) The [Gd-DTPA] which gives maximum SNR efficiency for each combination of TE and TR. This panel is used to determine the optimal concentration of Gd-DTPA for a given TE and TR. Right) SNR efficiency varying as a function of [Gd-DTPA], for fixed values of TE and TR. Four examples are shown, with TE/TR = i) 27/250 ms, ii) 27/800 ms, iii) 15/250 ms, iv) 15/800 ms, corresponding to points i - iv on panels A and B. The [Gd-DTPA] intervals resulting in 95% of the maximum SNR efficiency are indicated by shaded regions. The values shown in this figure were modelled using data from a region of interest in the corpus callosum in rats fixed with 2% PFA.

Collectively, these data suggest that the optimum SNR efficiency can be obtained by reducing fixative concentration, rehydrating samples for at least 17 days, selecting the shortest TE available, and choosing a TR and concentration of contrast agent to maximise SNR efficiency according to the results above. This strategy for optimisation is summarised in Table 3.

**Table 3:**
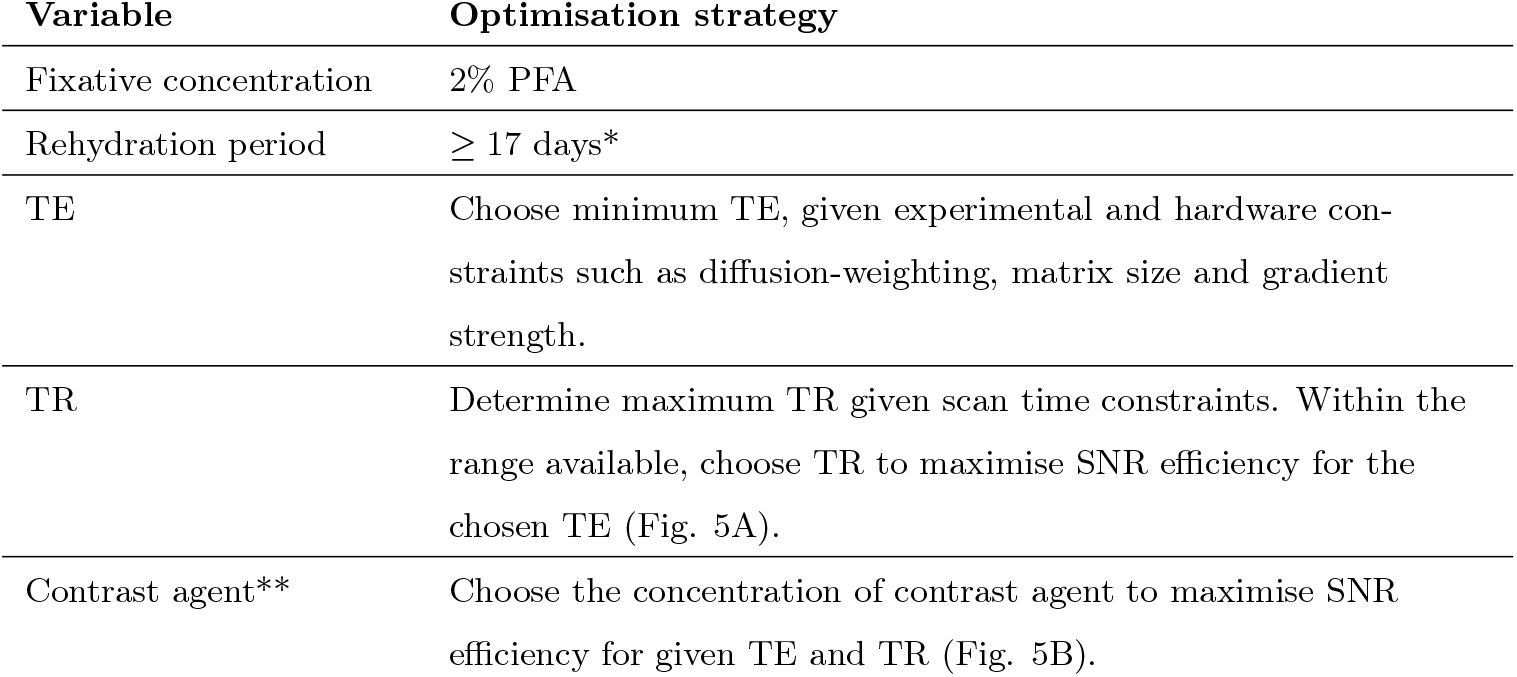
Optimisation strategy for tissue preparation and imaging parameters. ^*^This result was obtained from rehydration in a single volume of 50 ml PBS per sample. Increasing the volume or replacing the solution during this period may reduce the rehydration time further. ^**^In these experiments, the chosen concentration of Gd-DTPA contrast agent refers to the concentration in the perfusate. Samples were rehydrated in PBS with 1 mM Gd-DTPA.

### SNR comparison

To illustrate the improvement in SNR from using the above strategy, we compared acquisitions from rats prepared with and without tissue optimisation (Fig. 6). The standard sample, prepared without contrast agent and with the standard fixative concentration of 4% PFA, had SNR values of 12 and 10 in the cortex and white matter of the corpus callosum respectively in the non-diffusion-weighted image and 7 in both white matter and grey matter in a diffusion-weighted image with *b* = 2500 s/mm^2^. The optimised sample, fixed with 2% PFA and 15 mM Gd-DTPA, had SNR of 28 and 18 in grey and white matter respectively in the non-diffusion-weighted and 18 and 16 respectively in the diffusion-weighted volume. This represents an increase of SNR by a factor of 2.4 in grey and 1.9 in white matter in the non-diffusion-weighted and 2.6 and 2.2 for grey and white matter respectively in the in the diffusion-weighted images. A visible reduction in noise can be also directly seen in the optimised images. In terms of SNR efficiency, the optimised protocol resulted in an SNR per unit time (in minutes) of 9 in grey matter and 6 in white matter, compared with 4 and 3 in grey and white matter in the standard sample, for the non-diffusion-weighted volume. This represents an increase in SNR per minute by a factor of 2.4 and 1.9 in grey and white matter, respectively. The concentration of contrast agent for the optimised sample in this experiment was chosen to be within 95% of the maximum SNR efficiency for TE = 26.78 ms and TR 250 ms.

**Figure 6:**
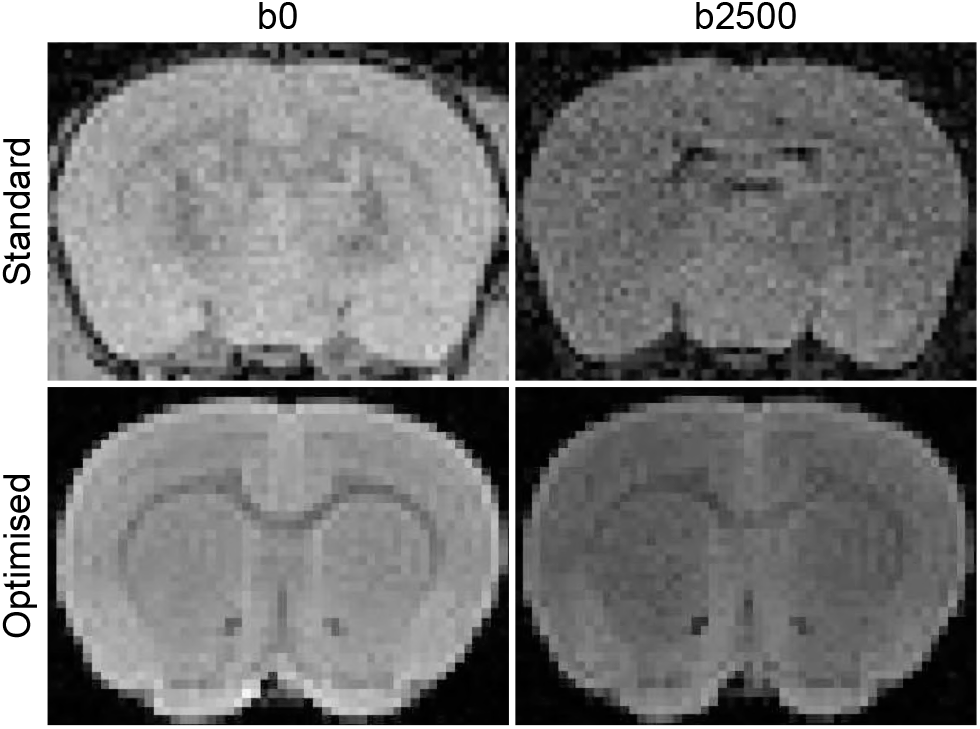
Improvement in SNR gained from optimising tissue and scanning parameters. The standard preparation included [Gd-DTPA] = 0mM, [PFA] = 4% (top row). The optimised tissue preparation was fixed with [Gd-DTPA] = 15mM, [PFA] = 2% (bottom row). Images are shown for non-diffusion-weighted (*b*0) and diffusion-weighted (*b*= 2500 s/mm^2^) images, using a 3D spin echo pulse sequence. The optimised sample was removed from the skull in preparation for subsequent high-resolution acquisition. Data from both samples were acquired using the same coil and protocol.

### High and mid-resolution diffusion imaging

One application for optimised tissue preparation and scanning parameters is to improve the quality of high-resolution MRI data. Here we present examples of diffusion datasets collected using the strategy outlined above. We include one example of a high-resolution diffusion weighted spin echo protocol and one mid-resolution diffusion weighted EPI protocol.

For the high-resolution acquisition, we chose a voxel size of 78 µm, a *b*-value of 4000 s/mm^2^ and 30 directions to ensure adequate diffusion contrast for tractography (Dyrby et al., 2011; Dell’Acqua & Tournier, 2019). We chose a maximum scan time of 96 hours to illustrate an example of a protocol that can practically be acquired over 4 days or a long weekend (the actual scan time was 91 hours 20 minutes). Given these constraints, we had a minimum TE of 26.77 ms and maximum TR of 250 ms. As per our strategy for maximising SNR efficiency, the sample was fixed with 2% PFA and 15 mM Gd-DTPA and rehydrated in PBS with 1 mM Gd-DTPA for at least 35 days. The resulting SNR of the non-diffusion weighted images before denoising was 12 in a cortical grey matter region and 9 in white matter measured at the midline of the corpus callosum. For comparison, we estimated that to achieve the same SNR with the standard sample preparation would require a TR of at least 750 ms, resulting in a tripling of the scan time to 11 days, 10 hours. This calculation assumes the same spatial resolution, field of view, TE and number of volumes, and is based on the parameters measured from samples fixed with 4% PFA, with no contrast agent, and 35 days rehydration in PBS (T1 = 1705 ms, T2 = 52 ms).

Figure 7 shows mean diffusion and non-diffusion-weighted images, DTI maps, and tractography reconstructions of the fornix, anterior commissure and corpus callosum. Of these tracts, the first two were dissected using DTI tractography, and illustrate the ability to resolve fine anatomical structures such as the body and the lateral fimbria of the fornix and the posterior arms of the anterior commissure. The corpus callosum was reconstructed using spherical deconvolution tractography because of its ability to resolve multiple fibre directions per voxel, for example where callosal and cortico-spinal fibres cross.

**Figure 7:**
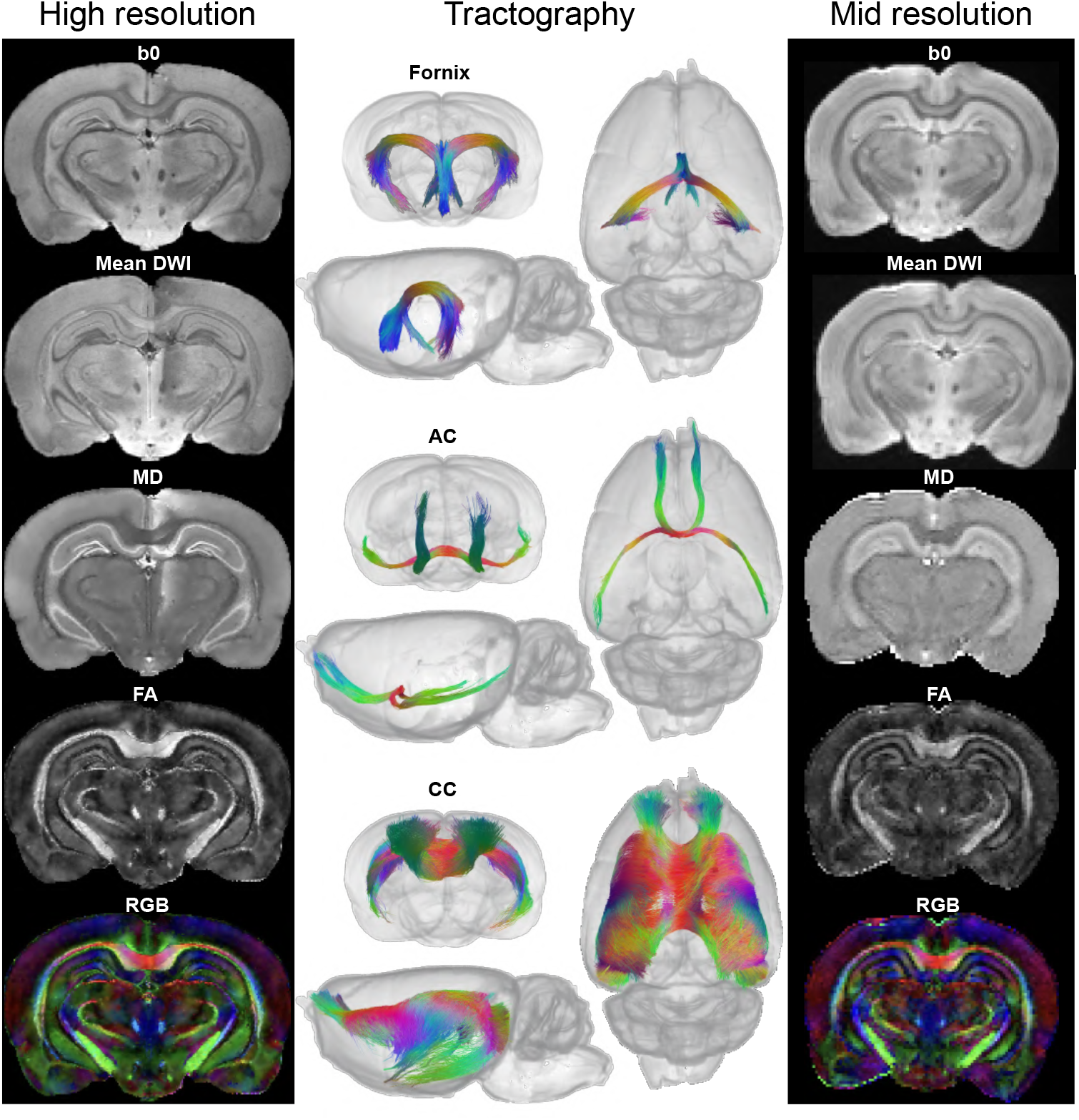
Diffusion data with optimised tissue preparation. Left) Diffusion maps for the high-resolution (78 µm) diffusion-weighted spin echo acquisition, including a non-diffusion-weighted image (b0), the mean of all diffusion-weighted imagines (Mean DWI), mean diffusivity (MD), fractional anisotropy (FA) and directional colour map with red-green-blue (RGB) encoding. Middle) Tractography reconstructions of the fornix and fimbria, anterior commissure (AC) and corpus callosum (CC) using the high-resolution data. Right) Diffusion maps for the mid-resolution (150 µm) diffusion-weighted EPI acquisition. Images represent data from a single rat brain

For applications where shorter scan time is prioritised over high-resolution, we also collected an example dataset of a diffusion weighted EPI protocol. In this example, we used the same tissue preparation and number of volumes as above: tissue perfused with 2% fixative and 15 mM Gd-DTPA, rehydrated in PBS with 1 mM Gd-DTPA, TE = 26.77ms, TR = 280ms, 4 averages and 30 diffusion directions, resulting in a scan time of 9 hours. We chose a *b*-value of 2000 s/mm^2^, which provides sufficient diffusion contrast if advanced diffusion modelling such as HARDI is not required, and a voxel size of 150 µm, sufficient to clearly resolve larger anatomical structures. The SNR in non-diffusion weighted images was 23 in grey and 17 in white matter. As SNR is proportional to the square root of the number of averages, this result shows that an adequate SNR for diffusion modelling, similar to that obtained in the high-resolution dataset, can be achieved in 2^1^/_4_ hours. Examples of diffusion and non-diffusion-weighted images and DTI maps are included in Figure 7.

Figure 8 shows, for example, that with this method, it is possible to distinguish between the cingulum and fibres fanning from the corpus callosum into the cortex, and to model the diffusion characteristics of the different layers of the hippocampus. These examples further illustrate the advantages and additional information that can be obtained at this resolution using HARDI modelling methods.**CC**

**Figure 8:**
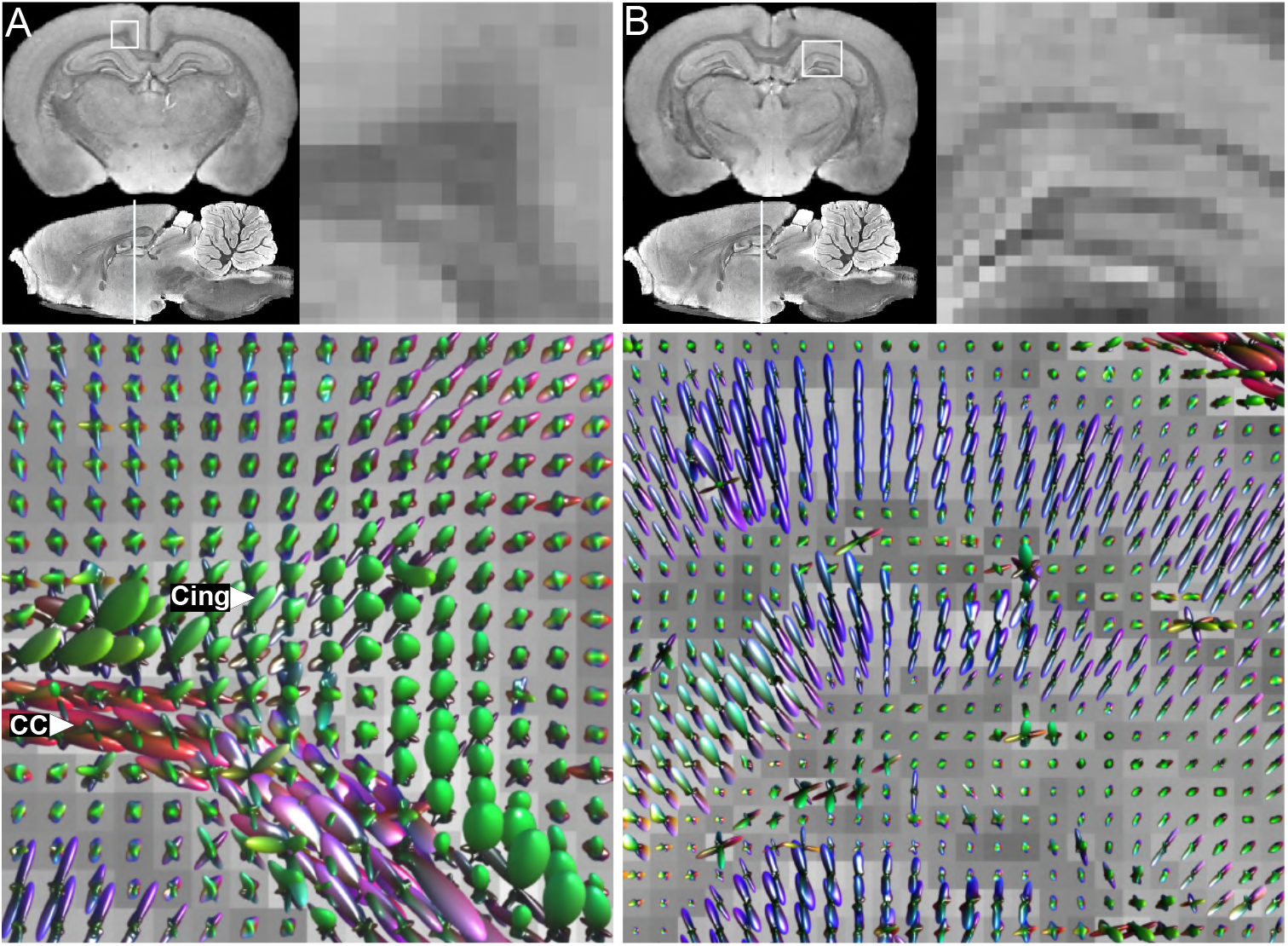
Fibre orientation distribution functions in high-resolution diffusion data. Regions of interest shown are A) a region including crossing fibres of the corpus callosum (CC) and cingulum (Cing.), and B) a region showing hippocampal layering. Top panels show region of interest location and enlarged view; bottom panels show fibre orientation distribution functions within the given region.

### Histology

Finally, we carried out histology to test whether there were any differences between the standard protocol (4% fixative, no contrast agent), the standard protocol with reduced fixative concentration (2% fixative, no contrast agent), and our optimised protocol (2% fixative with 15 mM Gd-DTPA). We found no differences in the quality of histology across the three samples, using Cresyl Violet to stain for Nissl bodies, IBA-1 for immuno- histochemistry staining of microglia or Luxol Fast Blue to stain for myelin. Figure 9 shows examples of these three stains applied to each sample, with inserts highlighting neurons and microglia in the hippocampus, and white matter in the internal capsule and striatum.

**Figure 9:**
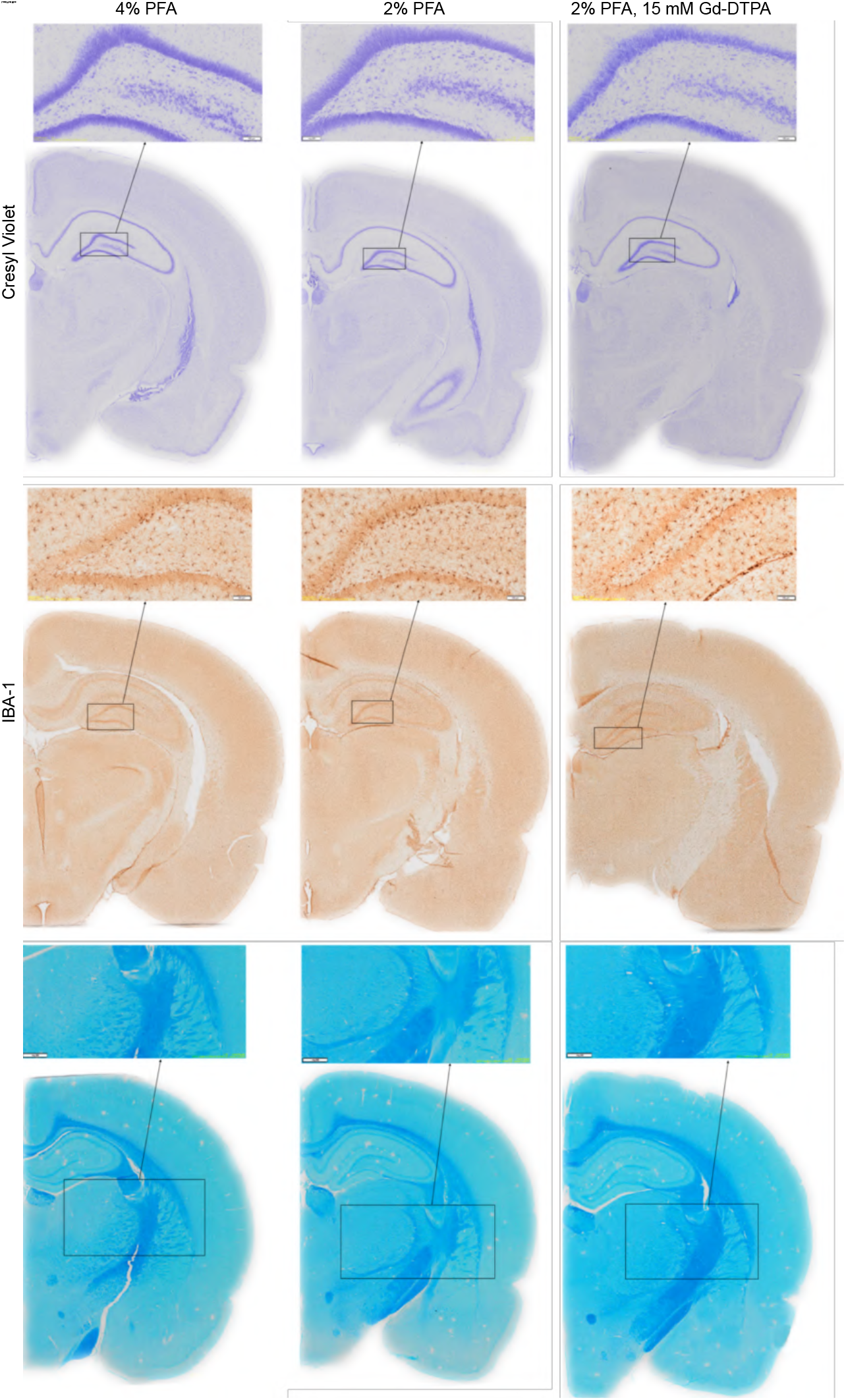
Histology showing no differences between different tissue preparations. The different preparations are compared in columns from left to right: 4% PFA; no contrast agent (standard protocol), 2% PFA, no contrast agent, 2% PFA, 15 mM Gd-DTPA added during perfusion (optimised protocol). Each sample was rehydrated in PBS, or PBS with 1 mM Gd-DTPA in the optimal case. From top to bottom the stains used are Cresyl Violet, IBA-1 antibody and Luxol Fast Blue.

## 4. Discussion

This study investigated how multiple tissue preparation factors can be combined to improve image quality in *ex vivo* diffusion MRI in the healthy adult male rat brain. We have shown how T1 and T2 can be manipulated together to maximise SNR efficiency by varying tissue rehydration time, fixative concentration and contrast agent concentration. We then presented an optimisation strategy for choosing these factors in combination with TE and TR. As an application of this approach, we included an example of a high quality, high-resolution diffusion protocol. The specific tissue preparation and scanning parameters used in this example may provide a valuable benchmark for future high-resolution *ex vivo* diffusion experiments, and our optimisation strategy in general may be used or adapted for a wide range of *ex vivo* MRI applications.

### Rehydration experiment

The aim of the rehydration experiment was to measure the relationship between rehydration time and tissue relaxation parameters to help determine how long rat brain samples should be soaked in PBS for before scanning. As in previous studies, we found that rehydration increased the T2 of fixed tissue, following an exponential recovery curve before plateauing, whereas it had comparatively little effect on T1 (Shepherd et al., 2005, 2009b; D’Arceuil et al., 2007; Leprince et al., 2015). In our results, the T2 in a corpus callosum region of interest from samples fixed with 4% and 2% PFA increased from 32 to 55 ms and 41 to 61 ms over the 35 day period measured. The results in grey matter measured in a cortical region of interest were similar, increasing from a T2 of 33 to 60 ms and 44 to 58 ms in the higher and lower fixative models, respectively. The rehydration-time curves for the grey and white matter regions were also similar.

The time required for each model to recover T2 to within 95% of its plateau value for the 4% and 2% fixative models respectively was 32 and 17 days in the corpus callosum region; and 47 and 14 days in the cortex. This shows that less rehydration time is needed for lower concentrations of fixative. It is worth noting that our results reflect the time required for the tissue to equilibrate in a single volume of 50 mL PBS solution at 4°C. Refrigeration was used here as a precaution against tissue degradation for samples fixed with reduced fixative. For comparison, previous studies with larger brain samples, standard fixative concentration and room-temperature rehydration with a single volume of PBS reported comparable time frames for T2 stabilisation compared to our 4% PFA model in the rat brain at 4°C. These include, for example, four to six weeks in the marmoset (Blezer et al., 2007) and three weeks in the ewe brain (Leprince et al., 2015). Other studies have reported much shorter equilibration times for small brain sections and/or frequent replacement of the PBS solution, for example, four days for 2cm-wide marmoset brain slices with a single PBS volume (D’Arceuil & de Crespigny, 2007), and 12 hours for 0.5 mm rat brain slices with four to five solution changes (Shepherd et al., 2009b). In our experiment, a single volume of PBS was used to provide a reference for how fast T1 and T2 stabilise in the most simple case, but the literature suggests that the T2 equilibration time could be further reduced in future experiments by increasing the volume or number of solution changes of PBS.

Shepherd et al. (2005, 2009b) have suggested that the reduction in T2 associated with formaldehyde fixation is due to factors including the presence of unbound fixative in the tissue and microstructural changes from the fixation reactions. The increase in T2 after soaking in PBS is suggested therefore to be due to unbound fixative being removed from the tissue. This could also explain why in our experiments, brains fixed with a lower concentration of fixative had higher T2, and stabilised faster while soaking in PBS.

The reduction in T1 of fixed tissue is thought to be caused by changes from chemical reactions that take place during fixation rather than the presence of free fixative in the tissue (Thelwall et al., 2006; Shepherd et al., 2005), which may explain why soaking samples in PBS does not cause a large increase in T1. In our results, the T1 of rats fixed with 4% and 2% PFA increased after 35 days from 1576 to 1618 ms and 1659 to 1781 ms respectively, in the corpus callosum region, and from 1631 to 1724 ms and 1742 to 1781 ms in the cortex region, again showing minimal difference between these two regions. While a longer T1 is generally disadvantageous for SNR efficiency, this was outweighed by the increase in SNR resulting from the prolongation of T2 as a result of rehydration. As in previous research (D’Arceuil et al., 2007), we found no substantial differences in FA and MD over time spent soaking in PBS. Rehydration is therefore an effective tool for manipulating T2 relaxation time to boost SNR.

### Gadolinium experiment

The addition of a gadolinium-based contrast agent was the next factor we investigated in order to alter T1 and T2 for improved SNR efficiency. We found that the relationship between observed relaxation times and the concentration of Gd-DTPA used in perfusion followed the theoretical model for contrast agent relaxivity (Lauffer, 1987) in line with previous *ex vivo* MRI studies (D’Arceuil et al., 2007; Ullmann et al., 2010). The rate of decline of T1 with respect to the concentration of Gd-DTPA was greater than that of T2, confirming that gadolinium-based contrast agents can be used to increase the T2 to T1 ratio, and therefore improve SNR efficiency. However, higher concentrations of contrast agent will eventually result in too much signal loss from T2 relaxation, hence the need for SNR optimisation in terms of [Gd-DTPA].

Our T2 data from rats fixed with 2% PFA deviated more from the predicted model compared with those fixed with 4% PFA. While the cause of this variation is not clear from our results, it is possible that fluctuations may be observed between rats acquired and perfused in different batches (Cahill et al., 2012). Nevertheless, we did not exclude these data points from the analysis, as doing so did not change the T1, T2 or SNR efficiency optimisation models substantially. Future studies may consider using rats acquired and perfused in the same batch where possible to minimise variability in T1 and T2 within an experiment. As for the rehydration experiment, the difference between the models based on measurements from white matter in the corpus callosum and grey matter in the cortex was minimal. This suggests that the same optimisation protocol can be applied effectively for both white and grey matter tissue.

Regarding the DTI measurements, we found no effect on FA or MD in our rat corpus callosum region of interest over the range of Gd-DTPA concentrations examined. In previous work by D’Arceuil et al. (2007), FA was also stable with respect to gadolinium concentration, whereas MD was lower overall and showed a decrease over concentrations of 0 to 10 mM Gd-DTPA. It is possible that factors such as the method of contrast agent staining (active vs. passive), the higher diffusion weighting used in D’Arceuil et al. (*b* = 4000 s/mm^2^) compared to our optimisation experiments, or potential differences in SNR might have influenced these differences in observed MD. The addition of Gd-DTPA is nonetheless an effective way to improve SNR efficiency while preserving sufficient diffusivity and anisotropy for fibre orientation modelling and tractography.

In addition to the relaxometry experiments above using Gd-DTPA, we also carried out measurements using the contrast agent gadobutrol for comparison, as at the time of writing gadobutrol is set to replace Gd-DTPA in some clinical settings. As expected from previous studies carried out in blood and plasma at lower field strengths, T1 and T2 relaxation times were shorter in tissue with gadobutrol than Gd-DTPA (Rohrer et al., 2005; Pintaske et al., 2006; Noebauer-Huhmann et al., 2010; Shen et al., 2015). We also observed that the ratio of T2 to T1 in the rat corpus callosum was higher for gadobutrol samples than for samples fixed with Gd-DTPA, suggesting that gadobutrol would produce even better SNR efficiency results than Gd-DTPA. The main analysis of this study focused on Gd-DTPA as the primary contrast agent in use and the most widely tested in previous preclinical studies, however future studies may consider using gadobutrol to further improve SNR efficiency, given the above results. Our data can be used to determine the amount of gadobutrol to be added during fixation to give the same T1 or T2 for a given concentration of Gd- DTPA, in fixed tissue prepared using a similar protocol to ours. This data will therefore be valuable for future studies in applying the proposed SNR efficiency optimisation strategy and comparing results with past literature based on Gd-DTPA.

The concentration values presented in our results refer to the amount of contrast agent added to the perfusate, which may differ from the true concentration of Gd-DTPA in the tissue at the time of scanning. Following previous studies, we used a higher concentration in the perfusion stage followed by a lower concentration in the rehydration stage, as the blood brain barrier inhibits some of the contrast agent from entering the tissue (Johnson et al., 2002, 2012; Calabrese & Johnson, 2013; Calabrese et al., 2013, 2015). The higher concentration is used because the blood-brain barrier inhibits the contrast agent from entering the tissue before fixation, whereas the tissue becomes more permeable to contrast agent after fixation (**?**). Due to both partial penetration of the contrast agent during fixation, and equilibration between the tissue and PBS solution during rehydration, we expect the true concentration to be lower than that which was added initially. Previous work by D’Arceuil et al. (2007), looking at the effect of contrast agent in the rehydration solution only, measured similar relaxivity values in the monkey brain at concentrations approximately one fifth of those we used in the perfusion. Rather than an indication of the concentration of gadolinium in the tissue, our results should be interpreted as a practical guide for how much contrast agent to use in the active staining protocol.

### Fixative concentration

Fixative concentration was another factor we examined to improve SNR efficiency. Throughout the experiments above, we examined the effect on relaxation times and diffusion metrics of lowering the concentration of fixative used from the standard 4% to 2%. In line with previous studies (Shepherd et al., 2009b; Birkl et al., 2018), we found that rats fixed with the lower concentration of fixative had longer T1 and T2 in the corpus callosum and required nearly half the time for rehydration compared to the standard 4% fixative model. However, the difference in relaxation times was less evident at high concentrations of Gd-DTPA. While the effect of changing the fixative concentration was relatively small compared to the effect of the contrast agent, our results suggest that it is a worthwhile factor to consider, in light of the shorter rehydration time required and the gains in T2 especially for concentrations of Gd-DTPA of ∼25 mM or less.

DTI metrics FA and MD were not affected by the change in fixative concentration in the rehydration experiment but in the gadolinium experiment FA appeared lower in the 2% fixative model. The mean FA was 0.33 in the 2% fixative model compared to 0.41 in the 4% fixative model. This could be due to changes in permeability between the two fixation models influencing the amount of gadolinium present in different compartments, however more investigation is required into the interaction of gadolinium and fixative concentration. These FA measurements were acquired with relatively low spatial resolution and low diffusion weighting, but the FA obtained from the 2% fixative model in the high-resolution dataset with a *b*-value of 4000 s/mm^2^ was 0.55 in an equivalent region of interest in the corpus callosum, and up to ∼0.7 in certain regions. These FA values are comparable to those reported in the literature for similar *ex vivo* diffusion acquisitions (D’Arceuil et al., 2007; Aggarwal et al., 2010; Dyrby et al., 2011; Calabrese et al., 2014), and were sufficient to produce high quality DTI maps and tractography reconstructions.

### SNR efficiency optimisation

Our strategy for SNR efficiency optimisation takes into account the effects of fixative concentration, tissue rehydration and contrast agent concentration as well as scanning parameters TE and TR. Following from the results discussed above, we first recommend using a reduced fixative concentration of 2% PFA and rehydrating the tissue until T2 stabilisation has occurred, to prolong T2 and maximise SNR efficiency. SNR efficiency can be further improved by perfusing the tissue with contrast agent at an optimal concentration determined in conjunction with TE and TR. In practice, TE is the parameter most constrained by factors such as diffusion weighting, acquisition matrix size, and gradient strength. TR may be constrained by the total scan time available and the number of diffusion directions required. Contrast agent, on the other hand, can be freely varied. We therefore propose that for a given TE, the TR and contrast agent concentration should be chosen together to maximise SNR efficiency, with an upper limit on TR based on scan time constraints if needed. This builds on the approach used in D’Arceuil et al. D’Arceuil et al. (2007), where the TR is chosen independently of the concentration of gadolinium. To our knowledge, D’Arceuil et al. D’Arceuil et al. (2007) is the only other study presenting a strategy to optimise gadolinium concentration quantitatively by modelling SNR efficiency, whereas other studies show a simple approach of comparing SNR or contrast to noise ratio between three or four given concentrations (Huang et al., 2009; Kim et al., 2009; Ullmann et al., 2010). Ours is also the only study we know of to include fixative as well as contrast agent concentration as variables and to present optimisation data for tissue fixed at 2% PFA.

In our example of tissue optimisation for a high-resolution diffusion protocol, we used 2% PFA, 15 mM Gd-DTPA, TE = 26.78 ms and TR = 250 ms. We found the SNR of this preparation to be more than double a baseline sample fixed with 4% PFA and no contrast agent, in a test acquisition including diffusion and non-diffusion weighted volumes. For comparison, other published examples of high-resolution *ex vivo* diffusion imaging with contrast enhancement have used TE/TR = 31.7/250 ms, [CA] in rehydration = 2 mM for the immersion-fixed macaque brain at 4.7 T (D’Arceuil et al., 2007); TE/TR = 21/100 ms, [CA] in fixation/rehydration = 5/2.5 mM for the immersion-fixed macaque brain at 7 T; TE/TR = 16.2/100 ms, [CA] in perfusion/rehydration = 50/5 mM for the perfusion-fixed rat brain at 7T (Johnson et al., 2012; Calabrese et al., 2013; Calabrese & Johnson, 2013); and TE/TR = 15/100 ms, [CA] in perfusion/rehydration = 50/5 mM for the perfusion-fixed mouse brain at 9.4 T (Calabrese et al., 2015). The optimal parameters for different studies may depend on various factors such as b-value, resolution, gradient strength, field strength and tissue relaxivity. For example, a lower b-value and lower resolution may be sufficient for certain applications, allowing a shorter TE, and leading to a shorter optimal TR and higher concentration of contrast agent. Scanners with lower gradient strength will generally necessitate a longer TE to achieve the same diffusion weighting, which would correspond to a longer optimal TR and lower concentration of contrast agent, according to our results in Figure 5. At higher field strengths, while more signal is available, T1 is generally longer (Stanisz et al., 2005) so we expect more gadolinium to be required to achieve maximum SNR efficiency. Our results may therefore be useful either as a benchmark for studies with similar experimental requirements and hardware, or as a reference point for future studies to undertaking tissue optimisation for SNR efficiency.

### High and mid-resolution diffusion imaging

Acquiring high-resolution *ex vivo* data is one application of our optimisation strategy. High-resolution diffusion imaging benefits from SNR efficiency optimisation because SNR is lost due to T2 relaxation when a long TE is required for diffusion encoding, and scan time is multiplied by the number of diffusion-weighted directions. As a result, *ex vivo* diffusion imaging studies are often limited in spatial resolution, number of diffusion directions and/or b-values.

Recently, other studies have used specialised hardware or advanced pulse sequences to address these challenges. For example, Johnson, Calabrese and colleagues achieved a voxel size of 50 µm in the rat brain using a custom-made radiofrequency transmit receive coil at 7T with 650 mT/m gradients for experiments with 6 directions and a *b*-value of approximately 1500 s/mm^2^ (Johnson et al., 2012; Calabrese et al., 2013; Calabrese & Johnson, 2013). Alternative pulse sequences have also been developed to reduce scan time without compromising SNR. Aggarwal et al. (2010), for example, used a diffusion-weighted gradient and spin echo sequence at 11.7 T with 3000 mT/m gradient strength, to achieve a voxel size of 55 µm in the mouse brain with 12 diffusion directions and a *b*-value of 1700 s/mm^2^. In comparison, using our tissue preparation and optimisation approach, at 9.4 T with a 1000 mT/m gradient set,, we were able to achieve a voxel size of 78 µm in the rat brain with 30 directions and a *b*-value of 4000 s/mm^2^. This study shows that with improved tissue preparation it is practical to achieve sufficient SNR, angular diffusion sampling and diffusion contrast for advanced modelling using the standard hardware and pulse sequences available on preclinical MRI systems.

The high-resolution diffusion data presented here was used to reconstruct fine anatomical details of white matter tracts with tractography. Applying spherical deconvolution methods allowed us to resolve crossing fibres in tracts such as the corpus callosum and visualise the fibre orientation distribution in cingulum area and hippocampal sublayers. We estimate that the same acquisition using the standard tissue preparation would require three times the scan time to achieve equivalent SNR. Applied in combination with improved pulse-sequences and customized hardware as mentioned above, the approach described here could further push the boundaries to even higher resolution diffusion acquisitions.

Improvements in SNR efficiency can equally be applied to low or mid-resolution acquisitions in order to improve data quality or decrease scan time. This could be applied for instance in studies where scanning time is limited, or large numbers of samples are required. As an example of a protocol that can be achieved in an overnight scanning session, we included a diffusion-weighted EPI acquisition with a voxel size of 150 µm, 30 directions, *b*= 2000 s/mm^2^, and scan time of 9.13 hours. This protocol included 4 signal averages, but the SNR measured was high enough that a single average would provide sufficient SNR for standard DTI measurements at a quarter of the time (2 hours, 17 minutes) if desired.

### Histology

Finally, we included histology data in this study to test whether tissue preparation optimised for MRI is compatible with histology or immunohistochemistry analysis. We compared samples fixed with 4% PFA, 2% PFA, and 2% PFA plus Gd-DTPA test if either reducing the fixative concentration or active staining had any detrimental effects to tissue fixation quality. We found no discernible differences between the three samples in the stains included, for cell bodies, microglia or myelin. To our knowledge, this is the first published example of histology data comparing 4% and 2% PFA-fixed tissue. A previous study by Cahill et al. (2012) found that MRI data was comparable from 2% and 4% PFA fixation in the mouse brain. This study also shows histology data from samples fixed with 4% with 0.5% PFA, the 0.5% PFA sample clearly showing tissue and cell degradation, which were not observed in our 2% PFA sample. Our data suggest that it is possible to achieve adequate fixation for histology using 2% PFA. Our histology results also showed no effect from the addition of 15 mM Gd-DTPA to the perfusate, in line with previous studies (Spencer et al., 2006; Lerch et al., 2011; Wang et al., 2019).

### Conclusion

In this study, we described the effects of tissue preparation factors on relaxivity and diffusivity of the corpus callosum in fixed rat brain tissue, and how these can be combined to optimise SNR efficiency. The approach we propose is to use a fixative concentration of 2% PFA, to rehydrate the tissue for at least 17 days, and to optimise the concentration of contrast agent depending on the minimum possible TE and corresponding optimal TR within one’s time constraints, using our results as a guide. The approach used here increased SNR by a factor of more than 2.5 compared to a standard preparation, and diffusion properties FA and MD were sufficiently preserved. We illustrated the application of this tissue optimisation approach with one high and one medium-resolution diffusion dataset, with examples of tractography with high-anatomical detail and multi-fibre modelling. In conclusion, our strategy will allow researchers to achieve faster acquisitions of high quality, high-resolution data for advanced diffusion analyses using a standard preclinical setup.

## Author contributions

**Rachel L. C. Barrett:** Conceptualisation, Methodology, Software, Formal analysis, Investigation, Writing - original draft, Visualisation, Project administration. **Diana Cash:** Resources, Writing - Review & Editing. **Camilla Simmons:** Resources. **Eugene Kim**: Resources, Writing – Review & Editing. **Tobias Wood:** Software, Writing – Review & Editing. **Richard Stones:** Writing – Review & Editing. **Anthony Vernon:** Writing – Review & Editing. **Marco Catani:** Funding acquisition. **Flavio Dell’Acqua:** Conceptualisation, Methodology, Software, Investigation, Writing - Review & Editing, Supervision.

## Acknowledgements

The authors thank Bernard Siow for contributing to our supply of Magnevist, Karen Randall for assisting with perfusion and tissue preparation, the members of the NatBrainLab for providing constructive feedback, and Steve Williams for facilitating our use of the preclinical MRI system. This research was supported by the Wellcome Trust [103759/Z/14/Z].

## References

Aggarwal, M., Mori, S., Shimogori, T., Blackshaw, S., & Zhang, J. (2010). Three-dimensional diffusion tensor microimaging for anatomical characterization of the mouse brain. Magn. Reson. Med., 64, 249–261. doi:10.1002/mrm.22426.

Alexander, D. C., & Barker, G. J. (2005). Optimal imaging parameters for fiber-orientation estimation in diffusion MRI. NeuroImage, 27, 357–367. doi:10.1016/j.neuroimage.2005.04.008.

Benveniste, H., & Blackband, S. J. (2002). Mr microscopy and high resolution small animal mri: applications in neuroscience research. Prog. Neurobiol., 67, 393–420. doi:10.1016/s0301-0082(02)00020-5.

Birkl, C., Soellradl, M., Toeglhofer, A. M., Krassnig, S., Leoni, M., Pirpamer, L., Vorauer, T., Krenn, H., Haybaeck, J., Fazekas, F., Ropele, S., & Langkammer, C. (2018). Effects of concentration and vendor specific composition of formalin on postmortem mri of the human brain. Magn. Reson. Med., 79, 1111–1115. doi:10.1002/mrm.26699.

Blamire, A. M., Rowe, J. G., Styles, P., & McDonald, B. (1999). Optimising imaging parameters for post mortem mr imaging of the human brain. Acta Radiol., 40, 593–597. doi:10.3109/02841859909175593.

Blezer, E. L. A., Bauer, J., Brok, H. P. M., Nicolay, K., & ‘t Hart, B. A. (2007). Quantitative mri-pathology correlations of brain white matter lesions developing in a non-human primate model of multiple sclerosis. NMR Biomed., 20, 90–103. doi:10.1002/nbm.1085.

Cahill, L. S., Laliberté, C. L., Ellegood, J., Spring, S., Gleave, J. A., van Eede, M. C., Lerch, J. P., & Henkelman, R. M. (2012). Preparation of fixed mouse brains for mri. NeuroImage, 60, 933–939. doi:10.1016/j.neuroimage.2012.01.100.

Calabrese, E., Badea, A., Cofer, G. P., Qi, Y., & Johnson, G. A. (2015). A diffusion mri tractography connectome of the mouse brain and comparison with neuronal tracer data. Cereb. Cortex, 25, 4628–4637. doi:10.1093/cercor/bhv121.

Calabrese, E., Badea, A., Watson, C., & Johnson, G. A. (2013). A quantitative magnetic resonance histology atlas of postnatal rat brain development with regional estimates of growth and variability. NeuroImage, 71, 196–206. doi:10.1016/j.neuroimage.2013.01.017.

Calabrese, E., Du, F., Garman, R. H., Johnson, G. A., Riccio, C., Tong, L. C., & Long, J. B. (2014). Diffusion tensor imaging reveals white matter injury in a rat model of repetitive blast-induced traumatic brain injury. Journal. Neurotrauma, 31, 938–950. doi:10.1089/neu.2013.3144.

Calabrese, E., & Johnson, G. A. (2013). Diffusion tensor magnetic resonance histology reveals microstructural changes in the developing rat brain. NeuroImage, 79, 329–339. doi:10.1016/j.neuroimage.2013.04.101.

D’Arceuil, H. E., & de Crespigny, A. J. (2007). The effects of brain tissue decomposition on diffusion tensor imaging and tractography. NeuroImage, 36, 64–68. doi:10.1016/j.neuroimage.2007.02.039.

D’Arceuil, H. E., Westmoreland, S., & De Crespigny, A. J. (2007). An approach to high resolution diffusion tensor imaging in fixed primate brain. NeuroImage, 35, 553–565. doi:10.1016/j.neuroimage.2006.12.028.

Dell’Acqua, F., Scifo, P., Rizzo, G., Catani, M., Simmons, A., Scotti, G., & Fazio, F. (2010). A modified damped richardson-lucy algorithm to reduce isotropic background effects in spherical deconvolution. NeuroImage, 49, 1446–1458. doi:10.1016/j.neuroimage.2009.09.033.

Dell’Acqua, F., & Tournier, J. D. (2019). Modelling white matter with spherical deconvolution: How and why? NMR Biomed., 32, e3945. doi:10.1002/nbm.3945.

Dyrby, T. B., Baaré, W. F. C., Alexander, D. C., Jelsing, J., Garde, E., & Søgaard, L. V. (2011). An ex vivo imaging pipeline for producing high-quality and high-resolution diffusion-weighted imaging datasets. Hum. Brain. Mapp., 32, 544–563. doi:10.1002/hbm.21043.

Dyrby, T. B., Innocenti, G. M., Bech, M., & Lundell, H. (2018). Validation strategies for the interpretation of microstructure imaging using diffusion mri. NeuroImage, 182, 62–79. doi:10.1016/j.neuroimage.2018.06.049.

Fishbein, K. W., Gluzband, Y. A., Kaku, M., Ambia Sobhan, H., Shapses, S. A., Yamauchi, M., & Spencer, R. G. (2007). Effects of formalin fixation and collagen cross-linking on t2 and magnetization transfer in bovine nasal cartilage. Magn. Reson. Med., 57, 1000–1011. doi:10.1002/mrm.21216.

Huang, S., Liu, C., Dai, G., Kim, Y. R., & Rosen, B. R. (2009). Manipulation of tissue contrast using contrast agents for enhanced MR microscopy in ex vivo mouse brain. NeuroImage, 46, 589–599. doi:10.1016/j.neuroimage.2009.02.027.

Johnson, G. A., Calabrese, E., Badea, A., Paxinos, G., & Watson, C. (2012). A multidimensional magnetic resonance histology atlas of the wistar rat brain. NeuroImage, 62, 1848–1856. doi:10.1016/j.neuroimage.2012.05.041.

Johnson, G. A., Cofer, G. P., Gewalt, S. L., & Hedlund, L. W. (2002). Morphologic phenotyping with mr microscopy: The visible mouse. Radiology, 222, 789–793. doi:10.1148/radiol.2223010531.

Jones, D. K. (2004). The Effect of Gradient Sampling Schemes on Measures Derived from Diffusion Tensor MRI: A Monte Carlo Study. Magnetic Resonance in Medicine, 51, 807–815. doi:10.1002/mrm.20033.

Jones, D. K., & Basser, P. J. (2004). “Squashing peanuts and smashing pumpkins”: How noise distorts diffusion-weighted MR data. Magn. Reson. Med., 52, 979–993. doi:10.1002/mrm.20283.

Kellner, E., Dhital, B., Kiselev, V. G., & Reisert, M. (2016). Gibbs-ringing artifact removal based on local subvoxel-shifts. Magnetic Resonance in Medicine, 76, 1574–1581. doi:10.1002/mrm.26054.

Kim, S., Pickup, S., Hsu, O., & Poptani, H. (2009). Enhanced delineation of white matter structures of the fixed mouse brain using Gd-DTPA in microscopic MRI. NMR in Biomedicine, 22, 303–309. doi:10.1002/nbm.1324.

Laitinen, T., Sierra, A., Bolkvadze, T., Pitkänen, A., & Gröhn, O. (2015). Diffusion tensor imaging detects chronic microstructural changes in white and gray matter after traumatic brain injury in rat. Frontiers in Neuroscience, 9. doi:10.3389/fnins.2015.00128.

Lauffer, R. B. (1987). Paramagnetic metal complexes as water proton relaxation agents for nmr imaging: theory and design. Chem. Rev., 87, 901–927. doi:10.1021/cr00081a003.

Leprince, Y., Schmitt, B., Chaillou, É., Destrieux, C., Barantin, L., Vignaud, A., Rivíere, D., & Poupon, C. (2015). Optimization of sample preparation for mri of formaldehydefixed brains. Proc. Int. Soc. Magn. Reson. Med., 23, 2283.

Lerch, J. P., Yiu, A. P., Martinez-Canabal, A., Pekar, T., Bohbot, V. D., Frankland, P. W., Henkelman, R. M., Josselyn, S. A., & Sled, J. G. (2011). Maze training in mice induces mri-detectable brain shape changes specific to the type of learning. NeuroImage, 54, 2086–2095. doi:10.1016/j.neuroimage.2010.09.086.

McNab, J. A., Jbabdi, S. S., Deoni, S. C. L., Douaud, G. G., Behrens, T. E. J., & Miller, K. L. (2009). High resolution diffusion-weighted imaging in fixed human brain using diffusion-weighted steady state free precession. Neuroimage, 46, 775–785. doi:10.1016/j.neuroimage.2009.01.008.

Miller, K. L., McNab, J. A., Jbabdi, S., & Douaud, G. G. (2012). Diffusion tractography of post-mortem human brains: optimization and comparison of spin echo and steady-state free precession techniques. Neuroimage, 59, 2284–2297. doi:10.1016/j.neuroimage.2011.09.054.

Miller, K. L., Stagg, C. J., Douaud, G., Jbabdi, S., Smith, S. M., Behrens, T. E., Jenkinson, M., Chance, S. A., Esiri, M. M., Voets, N. L., Jenkinson, N., Aziz, T. Z., Turner, M. R., Johansen-Berg, H., & McNab, J. A. (2011). Diffusion imaging of whole, post-mortem human brains on a clinical mri scanner. NeuroImage, 57, 167–181. doi:10.1016/j.neuroimage.2011.03.070.

Noebauer-Huhmann, I. M., Szomolanyi, P., Juras, V., Kraff, O., Ladd, M. E., & Trattnig, S. (2010). Gadolinium-based magnetic resonance contrast agents at 7 tesla: In vitro t1 relaxivities in human blood plasma. Invest. Radiol., 45, 554–558. doi:10.1097/RLI.0b013e3181ebd4e3.

Pfefferbaum, A., Sullivan, E. V., Adalsteinsson, E., Garrick, T., & Harper, C. (2004). Postmortem MR imaging of formalin-fixed human brain. NeuroImage, 21, 1585–1595. doi:10.1016/j.neuroimage.2003.11.024.

Pintaske, J., Martirosian, P., Graf, H., Erb, G., Lodemann, K. P., Claussen, C. D., & Schick, F. (2006). Relaxivity of gadopentetate dimeglumine (magnevist), gadobutrol (gadovist), and gadobenate dimeglumine (multihance) in human blood plasma at 0.2, 1.5, and 3 tesla. Invest. Radiol., 41, 213–221. doi:10.1097/01.rli.0000197668.44926.f7.

Roebroeck, A., Miller, K. L., & Aggarwal, M. (2019). Ex vivo diffusion mri of the human brain: Technical challenges and recent advances. NMR Biomed., 32, e3941. doi:10.1002/nbm.3941.

Rohrer, M., Bauer, H., Mintorovitch, J., Requardt, M., & Weinmann, H.-J. (2005). Comparison of magnetic properties of mri contrast media solutions at different magnetic field strengths. Invest. Radiol., 40, 715–24. doi:10.1097/01.rli.0000184756.66360.d3.

Shen, Y., Goerner, F. L., Snyder, C., Morelli, J. N., Hao, D., Hu, D., Li, X., & Runge, V. M. (2015). T1 relaxivities of gadolinium-based magnetic resonance contrast agents in human whole blood at 1.5, 3, and 7 t. Invest. Radiol., 50, 330–338. doi:10.1097/RLI.0000000000000132.

Shepherd, T. M., Flint, J. J., Thelwall, P. E., Stanisz, G. J., Mareci, T. H., Yachnis, A. T., & Blackband, S. J. (2009a). Postmortem interval alters the water relaxation and diffusion properties of rat nervous tissue {\textemdash} Implications for MRI studies of human autopsy samples. Neuroimage, 44, 820–826. doi:10.1016/j.neuroimage.2008.09.054.

Shepherd, T. M., Thelwall, P. E., Stanisz, G. J., & Blackband, S. J. (2005). Chemical fixation alters the water microenvironment in rat cortical brain slices - implications for mri contrast. Proc. Int. Soc. Magn. Reson. Med., 13, 619.

Shepherd, T. M., Thelwall, P. E., Stanisz, G. J., & Blackband, S. J. (2009b). Aldehyde fixative solutions alter the water relaxation and diffusion properties of nervous tissue. Magn. Reson. Med., 62, 26–34. doi:10.1002/mrm.21977.

Spencer, R. G., Fishbein, K. W., Cheng, A., & Mattson, M. P. (2006). Compatibility of gd-dtpa perfusion and histologic studies of the brain. Magn. Reson. Imaging, 24, 27–31. doi:10.1016/j.mri.2005.10.017.

Stanisz, G. J., Odrobina, E. E., Pun, J., Escaravage, M., Graham, S. J., Bronskill, M. J., & Henkelman, R. M. (2005). T1, T2 relaxation and magnetization transfer in tissue at 3T. Magnetic Resonance in Medicine, 54, 507–512. doi:10.1002/mrm.20605.

Sun, S.-W., Neil, J., Liang, H.-F., He, Y., Schmidt, R. E., Hsu, C. Y., & Song, S.-K. (2005). Formalin fixation alters water diffusion coefficient magnitude but not anisotropy in infarcted brain. Magn. Reson. Med., 53, 1447–1451. doi:10.1002/mrm.20488.

Sun, S.-W., Neil, J., & Song, S.-K. (2003). Relative indices of water diffusion anisotropy are equivalent in live and formalin-fixed mouse brains. Magn. Reson. Med., 50, 743–748. doi:10.1002/mrm.10605.

Thelwall, P. E., Shepherd, T. M., Stanisz, G. J., & Blackband, S. J. (2006). Effects of temperature and aldehyde fixation on tissue water diffusion properties, studied in an erythrocyte ghost tissue model. Magn. Reson. Med., 56, 282–289. doi:10.1002/mrm.20962.

Tournier, J. D., Calamante, F., & Connelly, A. (2013). Determination of the appropriate b value and number of gradient directions for high-angular-resolution diffusion-weighted imaging. NMR in Biomedicine, 26, 1775–1786. doi:10.1002/nbm.3017.

Tovi, M., & Ericsson, A. (1992). Measurements of T1 and T2 over time in formalin-fixed human whole-brain specimens. Acta Radiol., 33, 400–404. doi:10.1080/02841859209172021.

Ullmann, J. F. P., Cowin, G., Kurniawan, N. D., & Collin, S. P. (2010). Magnetic resonance histology of the adult zebrafish brain: optimization of fixation and gadolinium contrast enhancement. NMR Biomed., 23, 341–346. doi:10.1002/nbm.1465.

Veraart, J., Fieremans, E., & Novikov, D. S. (2016). Diffusion mri noise mapping using random matrix theory. Magnetic Resonance in Medicine, 76, 1582–1593. doi:10.1002/mrm.26059.

Wang, N., White, L. E., Qi, Y., Cofer, G., & Johnson, G. A. (2020). Cytoarchitecture of the mouse brain by high resolution diffusion magnetic resonance imaging. NeuroImage, 216, 116876. doi:10.1016/j.neuroimage.2020.116876.

Wang, N., Zhang, J., Cofer, G., Qi, Y., Anderson, R. J., White, L. E., & Johnson, G. A. (2019). Neurite orientation dispersion and density imaging of mouse brain microstructure. Brain Struct. Func., 224, 1797–1813. doi:10.1007/s00429-019-01877-x.

White, N. S., & Dale, A. M. (2009). Optimal diffusion MRI acquisition for fiber orientation density estimation: An analytic approach. Human Brain Mapping, 30, 3696–3703. doi:10.1002/hbm.20799.

Yong Hing, C. J., Obenaus, A., Stryker, R., Tong, K., & Sarty, G. E. (2005). Magnetic resonance imaging and mathematical modeling of progressive formalin fixation of the human brain. Magn. Reson. Med., 54, 324–332. doi:10.1002/mrm.20578.

